# Tra1 controls the transcriptional landscape of the aging cell

**DOI:** 10.1101/2022.07.13.499971

**Authors:** Khaleda Afrin Bari, Matthew D. Berg, Julie Genereaux, Christopher J. Brandl, Patrick Lajoie

## Abstract

Gene expression undergoes considerable changes during the aging process. The mechanisms regulating the transcriptional response to cellular aging remain poorly understood. Here, we employ the budding yeast *Saccharomyces cerevisiae* to better understand how organisms adapt their transcriptome to promote longevity. Chronological lifespan (CLS) assays in yeast measure the survival of non-dividing cells at stationary phase over time, providing insights into the aging process of post-mitotic cells. Tra1 is an essential component of both the yeast SAGA/SLIK and NuA4 complexes, where it recruits these complexes to acetylate histones at targeted promoters. Importantly, Tra1 regulates the transcriptional response to multiple stresses. To evaluate the role of Tra1 in chronological aging, we took advantage of a previously characterized mutant allele that carries mutations in the *TRA1* PI3K domain (*tra1*_*Q3*_). We found that loss of functions associated with *tra1*_*Q3*_ sensitized cells to growth media acidification and shortens lifespan. Transcriptional profiling reveals that genes differentially regulated by Tra1 during the aging process are enriched for components of the response to stress. Notably, expression of catalases (*CTA1, CTT1*) involved in hydrogen peroxide detoxification decreases in chronologically aged *tra1*_*Q3*_ cells. Consequently, they display increased sensitivity to oxidative stress. *tra1*_*Q3*_ cell*s* are unable to grow on glycerol indicating a defect in mitochondria function. Aged *tra1*_*Q3*_ cell*s* also display reduced expression of peroxisomal genes, exhibit decreased numbers of peroxisomes and cannot grow on media containing oleate. Thus, Tra1 emerges as an important regulator of longevity in yeast via multiple mechanisms.

## INTRODUCTION

*Saccharomyces cerevisiae* has been extensively studied as a eukaryotic model for lifespan regulation since many of the underlying molecular mechanisms are conserved from yeast to mammals ((Mortimer and Johnston 1959; Fontana *et al*. 2010; Kaeberlein 2010; Mirisola and Longo 2022). Chronological lifespan (CLS, defined as the extent of time non-dividing cells survive in a nutrient-deprived environment; (MacLean *et al*. 2001; Laun *et al*. 2006; Murakami *et al*. 2008) and replicative lifespan (the number of cell divisions before replicative senescence; (Steinkraus *et al*. 2008) are two distinct experimental aging models (Laun *et al*. 2006). Chronological lifespan experiments are carried out by culturing cells for a substantial period at the stationary phase. During chronological aging, yeast cells go through distinct growth phases: the mid-log phase, diauxic shift, and stationary phase. Each growth phase is illustrated by divergent metabolic activities and gene expression profiles, which are analogous to crucial characteristics of mammalian aging cells including cell cycle arrest, increased respiratory activity and lipid and protein homeostasis (Tissenbaum and Guarente 2002; Beach *et al*. 2013, 2015; Medkour and Titorenko 2016; Chadwick and Lajoie 2019; Chadwick *et al*. 2020; Mohammad *et al*. 2020). Studies of yeast chronological aging have enabled researchers to identify several key pathways that regulate longevity (Longo and Fabrizio 2012) such as Ras/Pka (Longo 1999) and Sch9/Tor (Fabrizio *et al*. 2001) and the role of sirtuins (Kaeberlein *et al*. 1999), not only in yeast but in other organisms.

How cells modulate their gene expression in response to stresses including aging involves all components of the transcriptional machinery. The recruitment of RNA polymerase II to promoters is tightly regulated by general transcription factors, gene-specific activators, and co-activators. Co-activators are multi-subunit protein complexes that interface between general transcription factors and gene specific activators (Kingston *et al*. 1996; Eberharter and Becker 2002; Helmlinger *et al*. 2011) and/or regulate modification of histone proteins and nucleosome remodeling (Roberts and Winston 1997; Rando and Winston 2012).Thus, co-activators play a vital role in global gene regulation. SAGA (Spt-Ada-Gcn5 acetyltransferase) and NuA4 (Nucleosome Acetyltransferase of H4) are prototypical multi-subunit co-activator protein complexes that are conserved among eukaryotic organisms. SAGA and NuA4 contain lysine acetyltransferases Gcn5 and Esa1, respectively, which acetylate both histone and non-histone proteins (Grant *et al*. 1997; Clarke *et al*. 1999; Steunou *et al*. 2014; Downey 2021). Transcriptome analyses for yeast strains lacking the SAGA components Gcn5 or Spt3 revealed that 10% of the stress-related genome is controlled by the SAGA complex (Huisinga and Pugh 2004). SAGA and NuA4 also participate in other non-transcriptional activities, such as telomere maintenance and DNA repair (Bird *et al*. 2002; Downs *et al*. 2004; Lin *et al*. 2008; Atanassov *et al*. 2009; Cheng *et al*. 2018, 2021). In addition, SAGA contains a deubiquitinating module (DUB) that includes the ubiquitin protease Ubp8 that cleaves ubiquitin from histone H2b (Henry *et al*. 2003; Morgan *et al*. 2016) and other targets. The additional targets for deubiquitination by the mammalian homolog of Ubp8, USP22, are particularly noteworthy for their role in cancer progression (Prokakis *et al*. 2021; Ning *et al*. 2022; De Luca *et al*. 2022).

Tra1 is uniquely found in both the SAGA and NuA4 complexes (Saleh *et al*. 1998; Grant *et al*. 1998; Allard *et al*. 1999). Tra1 belongs to the PIKK (phosphoinositide-3-kinase-related kinase) family, which includes Ataxia telangiectasia–mutated (ATM; Tel1 in *Saccharomyces cerevisiae*), ataxia telangiectasia and Rad3-related (ATR; Mec1 in *S. cerevisiae*), the DNA-dependent protein kinase catalytic subunit (DNA-PKc), mammalian target of rapamycin (mTOR; Tor1 and Tor2 in *Saccharomyces cerevisiae*) and SMG-1 (suppressor with morphological effect on genitalia family member) (Smith and Jackson 2003; Hill and Lee 2010; Shiloh and Ziv 2013). PIKK proteins have four common principal domains: an N-terminal HEAT (**<u>H</u>**untingtin, elongation factor 3 (**<u>E</u>**F3), Protein phosphatase 2A (PP2**<u>A</u>**), and **T**OR1), followed by FAT (FRAP-ATM-TRRAP), PI3K (phosphatidylinositol 3-kinase), and FATC (FRAP-ATM-TRRAP C-terminus) domains (Keith and Schreiber 1995); (Bosotti *et al*. 2000; Mordes *et al*. 2008). Approximately, half of the *TRA1* molecule consists of helical HEAT repeats and this section interacts with activator proteins to initiate transcription (Brown *et al*. 2001; Bhaumik *et al*. 2004; Knutson and Hahn 2011; Lin *et al*. 2012). A helical FAT domain located C-terminal to the HEAT repeats wraps the N-terminal section of the PI3K domain (Díaz-Santín *et al*. 2017; Sharov *et al*. 2017). At the C-terminus, PIKK family members share a highly conserved phosphoinositide-3-kinase (PI3K) domain. The PI3K domain consists of N- and C-terminal subdomains with a cleft between these two lobes, where ATP binds in the catalytically active PIKK family members (Huse and Kuriyan 2002; Taylor and Kornev 2011; Pavletich and Yang 2013). In all PIKK family members except Tra1/TRRAP, the PI3K domain regulates cell signaling by phosphorylating downstream target proteins. The PI3K domain of Tra1 lacks residues essential for ATP binding and phosphate transfer and therefore has no demonstrable kinase activity (Saleh *et al*. 1998; Nelson *et al*. 2006; Mutiu *et al*. 2007). Interestingly, however, mutation of conserved residues in the kinase cleft domain results in slow growth, increased stress sensitivity and decreased transcription of SAGA-regulated genes supporting a role for this pseudokinase domain (Berg *et al*. 2018).

In light of the importance of epigenetic changes in aging (Booth and Brunet 2016; Kane and Sinclair 2019), the involvement of SAGA and NuA4 in stress response processes and the unique role of Tra1 in both SAGA and NuA4 (Helmlinger *et al*. 2011; Cheung and Díaz-Santín 2019), we hypothesized that Tra1 contributes to longevity by regulating the transcriptional responses to stress associated with aging. Indeed, using a loss-of-function mutant, we find that Tra1 significantly alters the transcriptional landscape of the aging cell. Tra1 therefore emerges as a new regulator of the chronological aging process in yeast.

## MATERIALS AND METHODS

### Drugs

Oleic and myristic acid and diamide were purchased from Milipore-Sigma. Propidium iodide was from Thermo Fisher Scientific. 2-(*N*-morpholino)ethanesulfonic acid (MES) buffer solution was from Alfa Aesar.

### Strains and Plasmids

Yeast strains were constructed in the W303A background or BY4742 and are described in Table S1. Strains expressing either wild-type FLAG-tagged *TRA1* or *tra1*_*Q3*_ were created by integrating an *Sph*I-*Sac*I fragment from pCB2527 or pCB2537, respectively, into W303A and selecting *HIS+* cells expressing either wild-type *URA3*-FLAG-tagged *TRA1* or *tra1*_*Q3*_ PI3K domain and YCplac111-*DED1pr-YHR100* as previously described (Berg *et al*. 2018). yemRFP-SKL was generated by cloning yemRFP-SKL into the *Spe*I/*Sal*I sites of pRS416-GDP (Mumberg *et al*. 1995).

### Culture conditions and cell viability assay

Both the *TRA1* and *tra1*_*Q3*_ yeast cells (W303A derivatives) were grown to saturation overnight in synthetic complete medium (2% glucose with appropriate selection). An aliquot of the overnight cultures was diluted in fresh media and grown at 30°C in a rotating drum. Cell viability assays were conducted using propidium iodide as described previously (Chadwick *et al*. 2016). Briefly, cells were washed and resuspended in phosphate-buffered saline (PBS) containing 1 mM of propidium iodide. The positive control was prepared by boiling cells for 10 min before resuspending cells in propidium iodide staining solution. Unstained cells were used as a negative control. All samples were incubated for 10 min at room temperature in 96 well plates before imaging on a Gel doc system (Bio-Rad). The optical density (OD_600_) was measured using a BioTek Epoch 2 microplate reader. Both the *TRA1* and *tra1*_*Q3*_ cells were cultivated for several days and the cell viability of the chronologically aged cells was quantified at various time points throughout the aging process. Alternatively, the CLS assay was performed using caloric restricted medium (0.1% glucose), medium buffered with 0.1M MES buffer, or media containing .1% oleic acid, .1% myristic acid or 2% glycerol. To assess sensitivity to acetic acid, cells were grown in synthetic complete medium for four days and treated with acetic acid for 200 min with concentrations up to 0.08 mM and the cell viability assessed using propidium iodide. Survival rates were computed using the mean gray value from images along with OD_600_ using the ANALYSR program (Chadwick *et al*. 2016).

### Growth Assays

A single colony of yeast strain was inoculated and grown overnight in 2% synthetic medium without leucine at 30°C, 220 rpm. Overnight cultures were diluted at 1:10 fold ratio and OD_600_ measured using a spectrophotometer. Cell cultures were normalized to OD_600_ 0.1 and five-fold serial dilutions were spotted on solid media. Plates were incubated for 2 days at 30°C before taking images using a colony imager (S&P Robotics). Growth was quantified as previously described (Petropavlovskiy *et al*. 2020).

### RNA isolation and sequencing

Both *TRA1* and *tra1*_*Q3*_ cells were grown to saturation overnight, diluted 10-fold in 2% synthetic minus leucine medium and grown for 4 hours at 30°C in a rotating drum. An aliquot of the cells was spun down at day 0 and day 3 and stored at -80°C. Total RNA was isolated from cells using the RiboPure yeast kit (Thermo Fisher Scientific) according to the manufacturer’s instructions. RNA samples were prepared with three replicates for each genotype and time point. Quality of the RNA samples were analyzed using a Bioanalyzer to ensure a RIN value of 8 or higher.

Total RNA sequencing analysis was conducted by Azenta Life Sciences. RNA from each sample was converted into single stranded Illumina TruSeq cDNA libraries with poly dT enrichment. Libraries were sequenced on an Illumina HiSeq and each sample yielded between 27.5 and 39.7 million 150 bp paired-end sequencing reads. The raw reads and RNA count data were deposited in NCBI’s Gene Expression Omnibus (Edgar *et al*. 2002).

### Quality control, trimming, read alignment and differential gene expression analysis

Read quality was analyzed using FastQC (www.bioinformatics.babraham.ac.uk/projects/fastqc/). Trimmomatic default settings were used to trim low quality bases and adapter sequences (Bolger *et al*. 2014). Next, reads were aligned with the reference genome of *Saccharomyces cerevisiae S288C* sequence (R64-2-1; www.yeastgenome.org/) using STAR (Dobin *et al*. 2013). Only uniquely mapping reads were retained. featureCount was used to count the reads mapping to each gene (Liao *et al*. 2014). Differential expression analysis was performed with DESeq2 (Love *et al*. 2014). Differentially expressed genes with a Benjamini-Hochberg adjusted *P*-value cut-off of <= 0.05 were considered for further analysis.

Promoter analysis of differentially expressed genes (log2 fold change < 2 and*P*-value cut-off of <= 0.05) was performed using Yeastract (www.yeastract.com) (Monteiro *et al*. 2020).

### Fluorescence microscopy

*TRA1* and *tra1*_*Q3*_ cells expressing yemRFP-SKL were grown to saturation overnight, diluted 10 fold in 2% synthetic minus leucine medium and grown for 4 hrs at 30°C in a rotating drum. An aliquot of the cells was then spun down, washed, resuspended in PBS and transferred to a Nunc Lab-Tek chamber slide. Cells were imaged using a Zeiss Axiovert A1 wide-field fluorescence microscope with 63× 1.4 NA oil objective using a Texas Red filter (586 nm excitation/603 nm emission) and an AxioCam ICm1 R1 CCD camera. ImageJ was used to analyze the images (Schneider *et al*. 2012).

### Data availability

Strains and plasmids are available upon request. Gene expression data are available at GEO with the accession number: GSE206033.

## RESULTS

### Functional Tra1 is required for chronological aging

Tra1 is essential in yeast (Saleh et al., 1998) with the exception of the fission yeast *Schizosaccharomyce pombe* which expresses a second *TRA2* isoform (Helmlinger *et al*. 2011; Elías-Villalobos *et al*. 2019b). To determine the role of Tra1 in the aging process, we took advantage of a loss-of-function mutant that we previously characterized termed *tra1*_*Q3*_ (Berg *et al*. 2018). *tra1*_*Q3*_ contains three glutamine substitutions of arginine residues 3389, 3390 and 3456 proximal or within the putative ATP-binding cleft of the PI3K domain. *tra1*_*Q3*_ results in slow growth, increased stress sensitivity, transcriptional defects and impaired SAGA and NuA4 complex assembly (Berg *et al*. 2018; Razzaq *et al*. 2021). CLS was assessed by labeling cells with the viability dye propidium iodide (PI) at different time intervals during aging (**Figure 1a**). *tra1*_*Q3*_ cells display significantly reduced lifespan when compared to their wild-type counterparts (**Figure 1b,c,d**). Viability of *tra1*_*Q3*_ cells in exponentially growing culture is comparable to wild-type. *tra1*_*Q3*_ cells begin to decline in viability when cells reach the stationary phase indicating that a functional *TRA1* allele is required to extend CLS.

**Figure 1:**
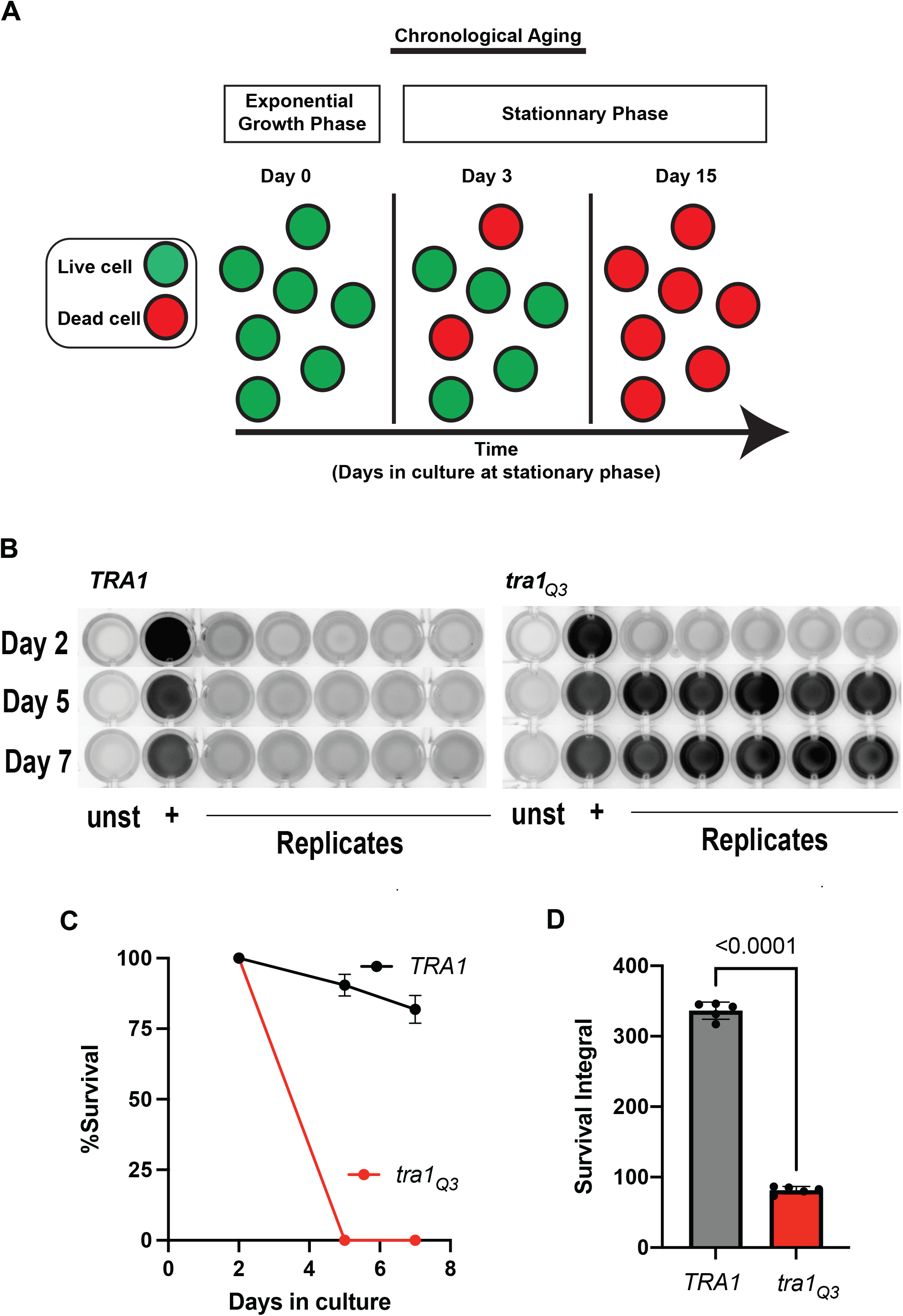
Functional Tra1 is required for chronological aging. **(A)** Chronological lifespan (CLS) is defined as the amount of time yeast cells survive at stationary phase. Experimentally, longevity is assessed by labeling cells with a viability dye such as propidium iodide at various time points during the aging process. Adapted from (Chadwick *et al*. 2016) **(B)** *tra1*_*Q3*_ cells display reduced CLS. *TRA1* and *tra1*_*Q3*_ cells were grown for the indicated times in standard synthetic complete medium and stained with propidium iodide to measure cell survival. Stained cells were imaged in a 96 well plate. For each time point, a negative control (unstained cells), a positive control (boiled cells) and 5 replicates were analyzed. **(C)** Normalized survival rates over the aging process and **(D)** survival integral are shown in the bar graph. Significance was assessed using an unpaired Student T-test.

Since caloric restriction efficiently increases longevity in yeast and other organisms (Jiang *et al*. 2000; Fontana *et al*. 2010; Choi *et al*. 2013; Arlia-Ciommo *et al*. 2014; Leonov *et al*. 2017; Campos *et al*. 2018), we then tested whether functional Tra1 is required for lifespan extension by caloric restriction. Caloric restriction is achieved by growing the cells in 0.1% glucose medium whereas complete medium contains 2% glucose. We found that when grown under caloric restriction conditions, *tra1*_*Q3*_ cells display extended lifespan similar to wild-type *TRA1* cells (**Figure 2**) suggesting that Tra1 function is dispensable for lifespan extension by caloric restriction.

**Figure 2:**
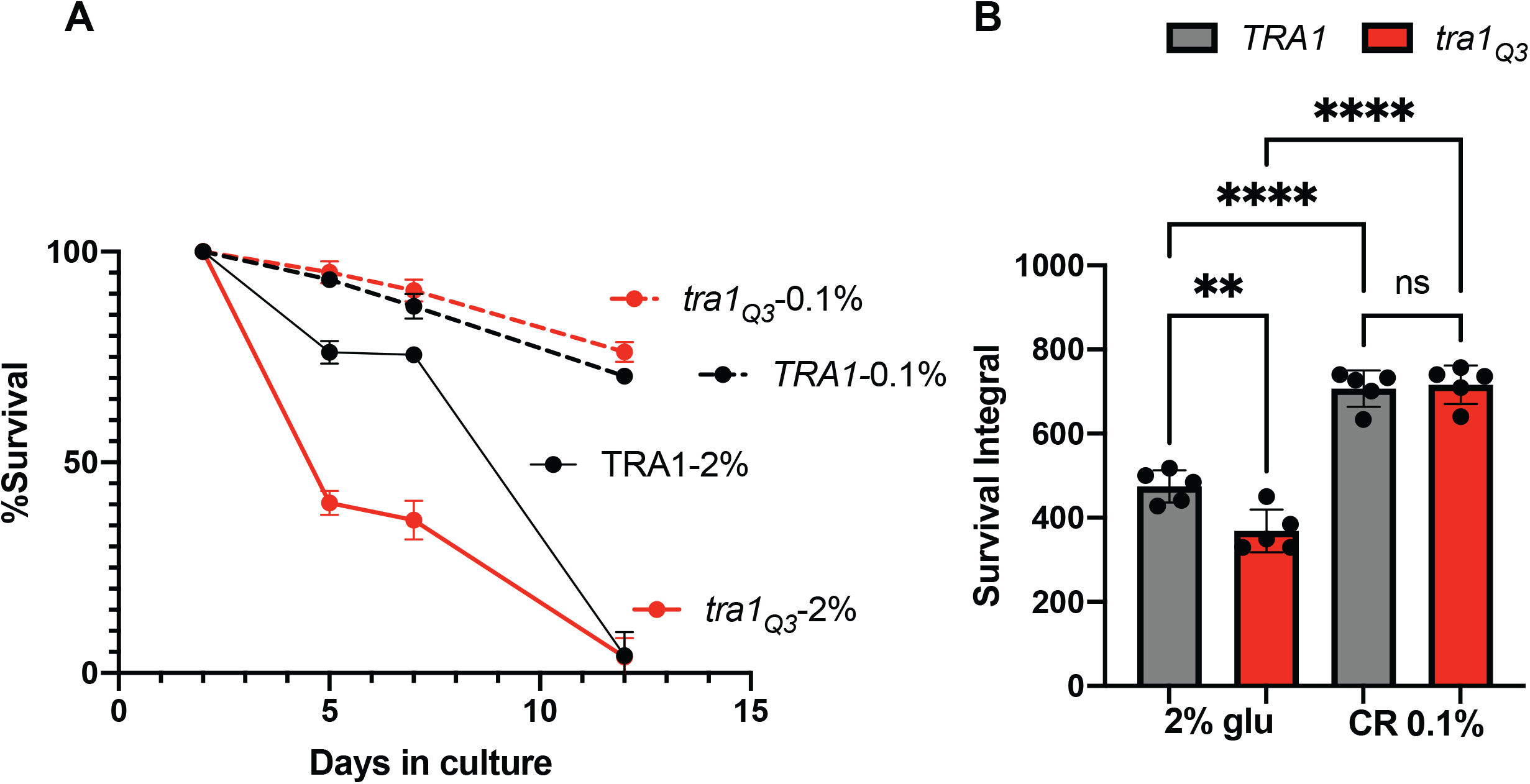
Tra1 function is dispensable for lifespan extension by caloric restriction. **(A)** *TRA1* and *tra1*_*Q3*_ cells were grown for the indicated time in standard synthetic complete medium containing either 2% glucose or 0.1% glucose (CR) and stained with propidium iodide to measure cell survival. Stained cells were imaged in a 96 well plate. For each time point, a negative control (unstained cells), a positive control (boiled cells) and 5 replicates were analyzed. **(B)** Normalized survival rates over the aging process and calculated survival integral are shown in graphs. Significance was assessed using a one way anova followed by a Tukey multiple comparisons test. **p<0.01 ***p<0.005 ****p<0.0001

### *tra1*_*Q3*_ cells are more sensitive to acetic acid

Culture media acidification is a predominant cell-extrinsic factor associated with cell death during chronological aging (Burtner *et al*. 2009; Burhans and Weinberger 2009; Hu *et al*. 2014; Deprez *et al*. 2018; Chaves *et al*. 2021). During chronological aging, acetic acid is produced following depletion of glucose from the growth media when ethanol is utilized as the main carbon source. Indeed, long lived mutants such as *sch9Δ* and *ade4Δ* show increased resistance to acetic acid (Burtner *et al*. 2009; Matecic *et al*. 2010). Therefore, we sought to determine if increased sensitivity to acidification of the cell culture media by acetic acid explains, at least in part, the shorter lifespan of *tra1*_*Q3*_ cells. To this end, *TRA1* and *tra1*_*Q3*_ cells were grown to stationary phase for 4 days and treated with various concentrations of acetic acid. Indeed, *tra1*_*Q3*_ increases sensitivity to acetic acid compared with wild-type *TRA1* (**Figure 3a,b**). Media acidification during chronological aging is alleviated by buffering the cultures to pH 6 with citrate phosphate or 2-(*N*-morpholino)ethanesulfonic acid (MES) resulting in increased longevity (Burtner *et al*. 2009; Burhans and Weinberger 2009). We found that buffering the aging media pH with MES significantly extended the lifespan of both *TRA1* and *tra1*_*Q3*_ cells (**Figure 3c,d**), suggesting the inability to respond to acid stress plays an important role in the reduced longevity phenotype associated with the *tra1*_*Q3*_ mutation.

**Figure 3:**
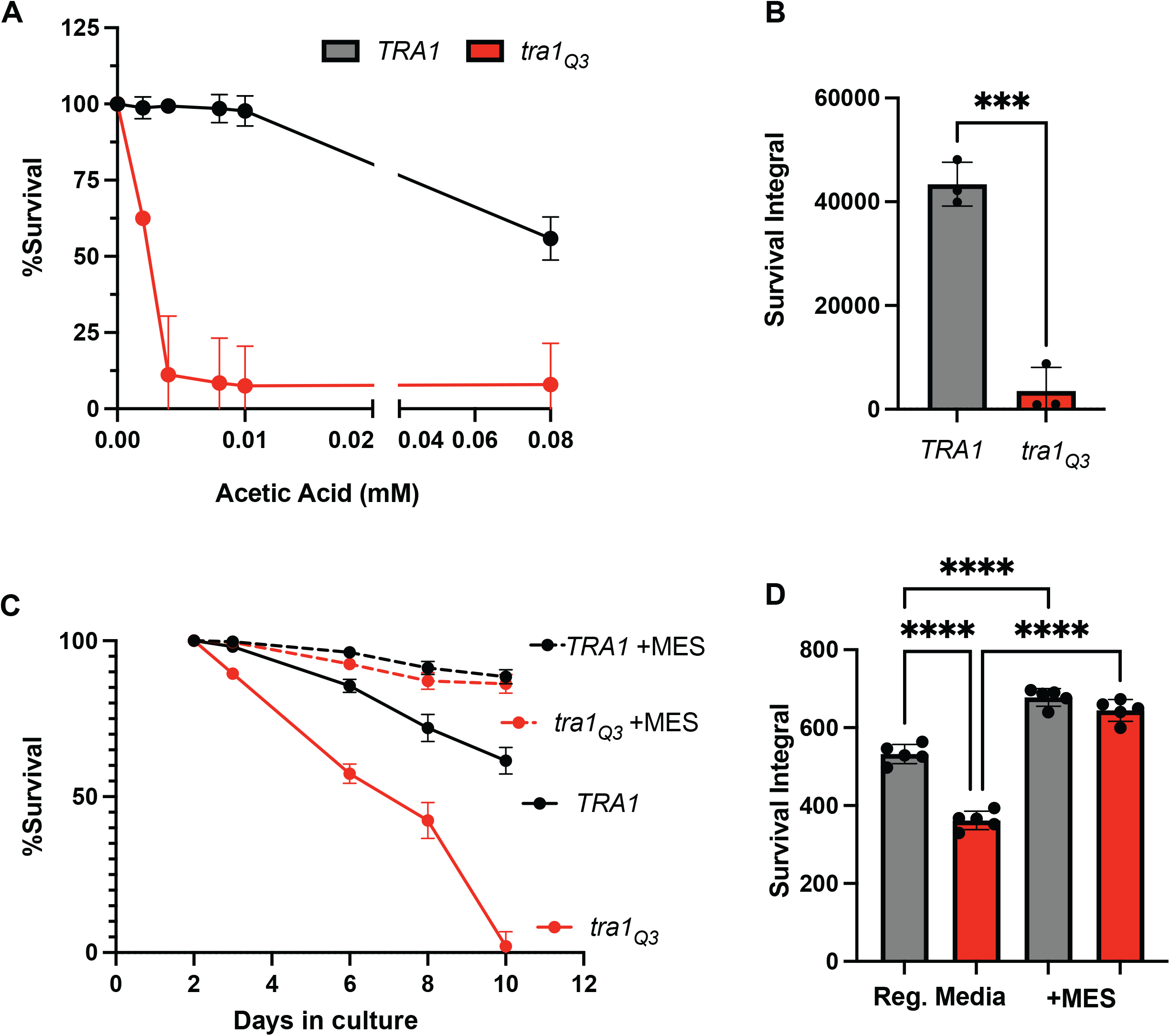
Increased sensitivity of *tra1*_*Q3*_ cells to media acidification is linked to shortened lifespan. **(A)** *tra1*_*Q3*_ cells display increased sensitivity to acetic acid. *TRA1* and *tra1*_*Q3*_ cells were grown in the presence of the indicated concentrations of acetic acid and stained with propidium iodide to measure cell survival over the aging process. For each time point, a negative control (unstained cells), a positive control (boiled cells) and 5 replicates were analyzed. **(B)** Normalized survival rates upon acetic acid treatment were calculated and survival integrals are shown. **(C)** *TRA1* and *tra1*_*Q3*_ cells were grown for the indicated time in standard synthetic complete medium with or without 0.1 M MES and stained with propidium iodide to measure cell survival. Stained cells were imaged in a 96 well plate. For each time point, a negative control (unstained cells), a positive control (boiled cells) and 5 replicates were analyzed. **(D)** Normalized survival rates over the aging process and calculated survival integrals are shown. Significance was assessed using a one way anova followed by a Tukey multiple comparisons test. ***p<0.005 ****p<0.0001

### Tra1 controls the transcriptional landscape of the aging cell

We previously found that Tra1 regulates the expression of genes associated with several stress responses including cell wall stress and the response to misfolded proteins (Berg *et al*. 2018; Jiang *et al*. 2019; Razzaq *et al*. 2021). Based on findings that *tra1*_*Q3*_ cells have a shorter lifespan, we next sought to determine how the loss of Tra1 function impacts the transcriptional landscape of the aging cell. We performed a global transcriptome analysis using RNA sequencing for *TRA1* and *tra1*_*Q3*_ cells before chronological aging (day 0) and after three days at stationary phase (day 3) to identify genes subjected to different transcriptional regulation. We found that ∼60% of the variance in the transcriptome data is linked to the aging process and ∼20% is linked to the *tra1*_*Q3*_ mutation (**Figure 4a**). The larger distance between *TRA1* and *tra1*_*Q3*_ cells after aging suggests that transcriptional differences between the two strains is greater in aged cells. At day 0, 394 genes were statistically differentially regulated (175 downregulated and 219 upregulated) when comparing *TRA1* and *tra1*_*Q3*_ cells (p < 0.05, log_2_ fold change > 2) (Table S2). At day 3, there were 2026 statistically differentially expressed genes (978 downregulated and 1048 upregulated)(**Table S2**). When comparing the age-genotype interaction scores (Smith and Kruglyak 2008; Sardi and Gasch 2018), we found 287 genes (161 negative and 126 positive) with a log_2_ score > 2.5 and p < 0.05, suggesting that these genes differentially respond to the aging process (**Figure 4b**) in the two genotypes (*TRA1* and *tra1*_*Q3*_). Gene Ontology analysis revealed significant enrichment in several biological processes, including RNA binding, transmembrane transporter activity, oxidoreductase activity and unfolded protein binding (**Figure 4c**). Genes associated with misfolded protein stress (*SSA1, HSP26, HSP42, SSA4, MID1, KAR2, SSA2, CPR6, HSP60, HSC82, APJ1, HSP10, HSP82)* are upregulated in aged *tra1*_*Q3*_ cells compared to wild-type suggesting that these cells exhibit increased proteotoxic stress. Genes with a negative age-genotype interaction score were also enriched in Rlm1 target genes, consistent with our previous results showing that Tra1 controls cell wall integrity (**Figure S1**) (Berg *et al*. 2018; Razzaq *et al*. 2021). Genes with the highest positive and negative interaction scores are presented in **Figure 4d**. Genes upregulated in aged *tra1*_*Q3*_ cells compared to wild-type include *RGL1* (a regulator of Rho1 signaling) and *BTN2* (a v-snare binding protein). Interestingly, *BTN2* expression is increased under severe ethanol stress (Yamauchi and Izawa 2016), reinforcing the idea that *tra1*_*Q3*_ cells are subject to increased stress during the aging process. Genes with positive age-genotype interaction are upregulated in wild-type *TRA1* compared to *tra1*_*Q3*_ cells in aged cells. Examples of such genes are *SIP18* and *MLS1. SIP18* encodes a phospholipid binding hydrophilin involved in vacuolar membrane fusion and is upregulated during replicative lifespan (Ghavidel *et al*. 2018). *MLS1* encodes a malate synthase involved in the utilization of non-fermentable carbon sources and is also upregulated in replicatively aged cells (Lesur and Campbell 2004).

**Figure 4:**
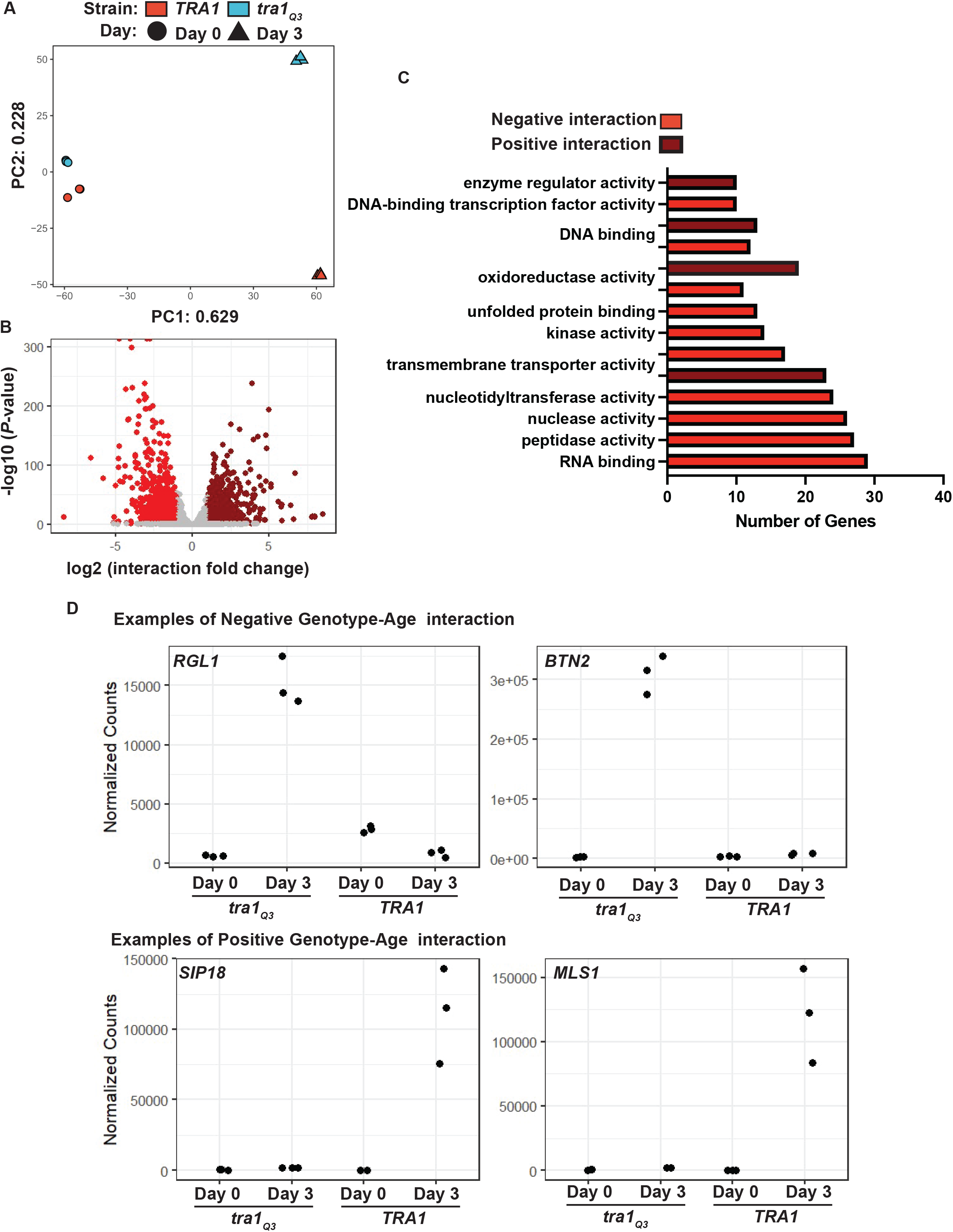
The *tra1*_*Q3*_ mutation alters the transcriptional landscape of aging cells. **(A)** Principal component analysis of centered log ratio of normalized reads from *TRA1* and *tra1*_*Q3*_ cells at day 0 and day 3. Each point represents a single biological replicate (n=3). **(B)** Volcano plot of genes that respond differently to the aging process in the *tra1*_*Q3*_ cells compared to wild-type *TRA1* (coloured points represent p < 0.05, log_2_ fold change > 1). **(C)** Significantly enriched GO biological processes were determined for genes with both positive and negative age-genotype interaction with log_2_ interaction score > 2 (p < 0.05) in *tra1*_*Q3*_ cells compared to wild-type. **(D)** Examples of genes with positive (*SIP18, MLS1*) and negative (*RGL1, BTN2*) age-genotype interaction. Normalized RNA sequencing read counts are shown for *TRA1* and *tra1*_*Q3*_ cells at day 0 and day 3.

We previously showed that cells expressing *tra1*_*Q3*_ increase expression of *TRA1* as a possible compensatory mechanism for the loss of protein function (Berg *et al*. 2018). Here, we observed this phenomenon as *tra1*_*Q3*_ mRNA increased ∼1.7 fold compared to wild-type at day 0. This difference was exacerbated in aged cells as *TRA1* mRNA increased ∼3.2 fold in *tra1*_*Q3*_ cells after aging (**Figure 5a**). We also analyzed the mRNA levels of other SAGA and NuA4 components (**Figure 5b**). Most striking are the upregulation of *ADA2* in aged *tra1*_*Q3*_ cells and downregulation of components of the SAGA DUB module (*UBP8, SUS1, SGF73, SGF11*). Among the NuA4 components, *ARP4* was upregulated in aged *tra1*_*Q3*_ cells. Interestingly, mutation of *ARP4* affects chronological lifespan in combination with deletion of the linker histone Hho1 (Vasileva *et al*. 2021).

**Figure 5:**
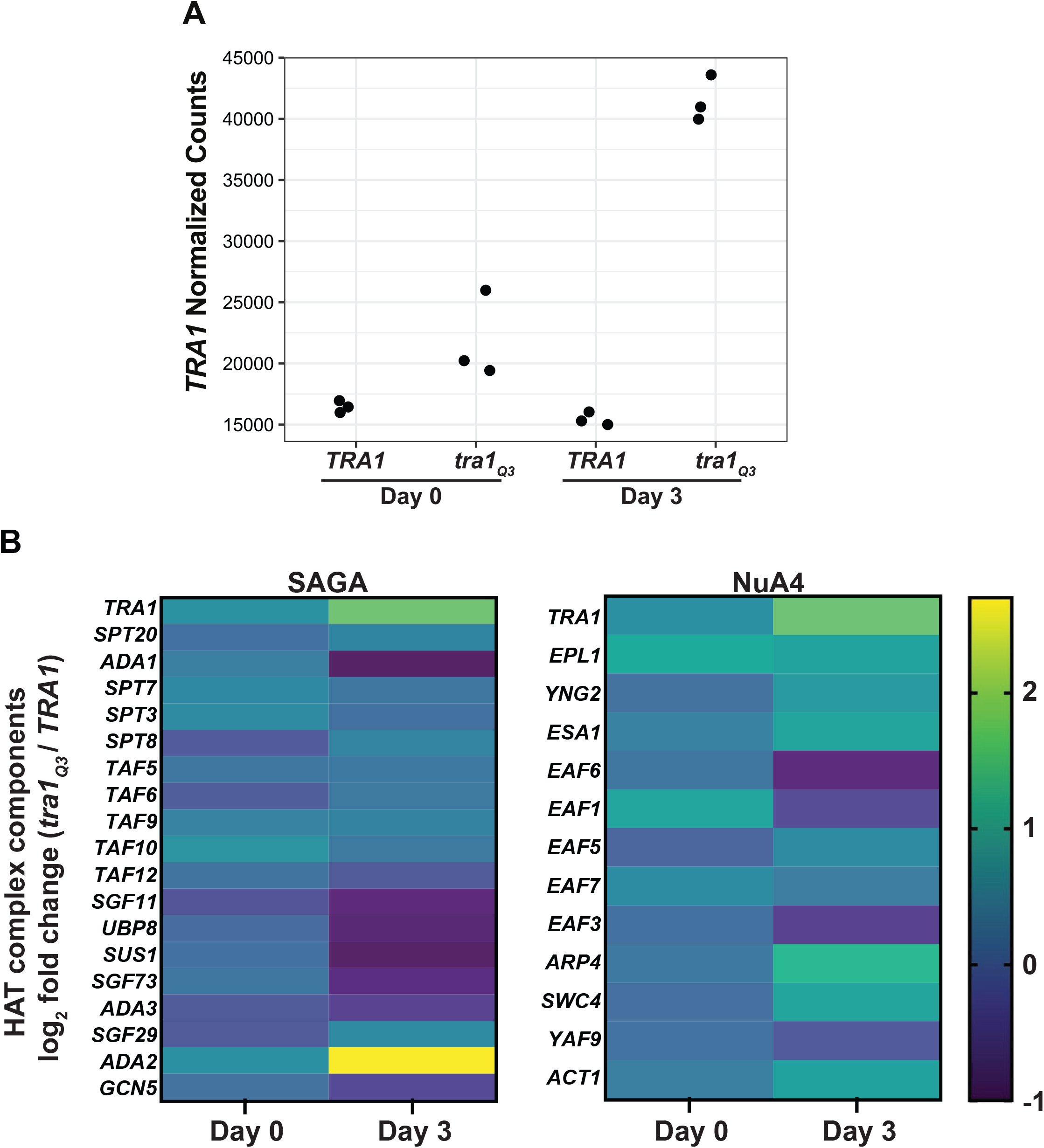
Aged *tra1*_*Q3*_ cells display increased *TRA1* mRNA abundance. **(A)** Normalized RNA sequencing read counts are shown for *TRA1* mRNA in wild-type *TRA1* and *tra1*_*Q3*_ cells at day 0 and day 3. **(B)** log_2_ fold change (*tra1*_*Q3*_*/TRA1*) for the mRNA of genes encoding SAGA and NuA4 components at day 0 and day 3.

GO analysis of genes that respond differentially to the aging processing in *tra1*_*Q3*_ cells revealed an enrichment in genes associated with oxidoreductase activity (**Figure 6a**). Among those genes were *CTT1* and *CTA1*, which encode two versions of catalase in *S. cerevisiae*. Ctt1 is cytoplasmic (Seah and Kaplan 1973) while Cta1 localizes to both the mitochondria and peroxisomes (Petrova *et al*. 2004). Catalase activity is crucial for hydrogen peroxide detoxification and is an important regulator of oxidative stress resistance associated with various conditions, including aging in yeast (Petriv and Rachubinski 2004; Agarwal *et al*. 2005; Mesquita *et al*. 2010; Rona *et al*. 2015; Guaragnella *et al*. 2019). Both genes are substantially upregulated in aged wild-type *TRA1* cells but not in *tra1*_*Q3*_ cells (**Figure 6b**). Consequently, *tra1*_*Q3*_ cells were more sensitive to oxidative stress induced by diamide (**Figure 6c**). Thus, these data indicate that *tra1*_*Q3*_ cells have reduced capacity to cope with oxidative damage that is usually associated with the aging process (Pan 2011).

**Figure 6:**
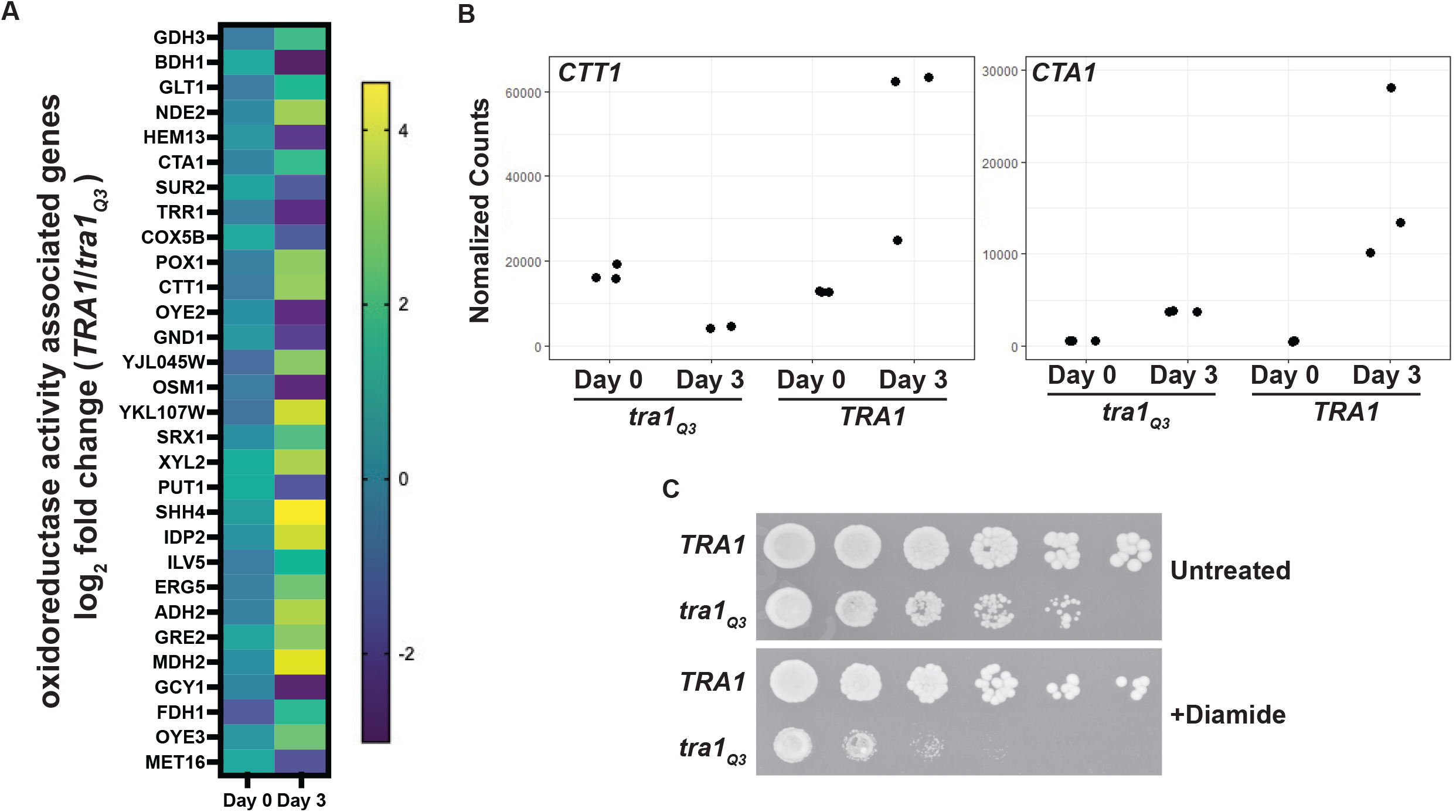
*tra1*_*Q3*_ cells are sensitive to oxidative stress. **(A)** log_2_ fold change (*TRA1/tra1*_*Q3*_) for mRNA of genes associated with oxidoreductase activity in wild-type *TRA1* and *tra1*_*Q3*_ cells at day 0 and day 3. **(B)** Normalized RNA sequencing read counts are shown for *CTT1* and *CTA1* in *TRA1* and *tra1*_*Q3*_ cells at day 0 and day 3. **(C)** *tra1*_*Q3*_ cells are sensitive to diamide. *TRA1* and *tra1*_*Q3*_ cells were spotted onto agar plates without (untreated) or with 1 mM diamide.

Several differentially expressed genes were also linked to peroxisomal **β**-oxidation and fatty acid metabolism (*POT1, POX1, MLS1, DCI1, ECI1, TES1*) (**Figure 7a**). The **β**-oxidation pathway and peroxisome proliferation are crucial for chronological aging (Lefevre *et al*. 2013). In yeast, **β**-oxidation solely occurs in peroxisomes (Hiltunen *et al*. 2003) and strains with defective peroxisomes fail to grow in presence of oleate due to their incapacity to use fatty acid as a carbon source (Lockshon *et al*. 2007). We found that *tra1*_*Q3*_ cells were unable to grow on media containing oleate or myristate, suggesting that they are defective in **β**-oxidation (**Figure 7b**). Because acetyl-CoA produced by **β**-oxidation serves as an energy source only in respiratory-competent strains, we assessed growth on glycerol. We found that *tra1*_*Q3*_ cells are unable to grow on plates containing glycerol (Figure 7b). Since **β**-oxidation in yeast is performed solely in peroxisomes, we also analyzed the expression of *PEX34* and *PEX21* in *TRA1* and *tra1*_*Q3*_ cells during the aging process. Pex34 regulates peroxisome biogenesis (Tower *et al*. 2011). Pex21 regulates import of protein into the peroxisomal matrix (Purdue *et al*. 1998). Aged *tra1*_*Q3*_ cells show reduced expression of *PEX34* and *PEX21* compared to aged wild-type cells (**Figure 7c**). *tra1*_*Q3*_ cells consequently display a reduced number of peroxisomes when labeled with the fluorescent reporter yemRFP-SKL at both day 0 and day 3 (**Figure 7d,e**). Therefore, our data suggest that *tra1*_*Q3*_ cells have defective peroxisome function that might contribute to their reduced chronological lifespan.

**Figure 7:**
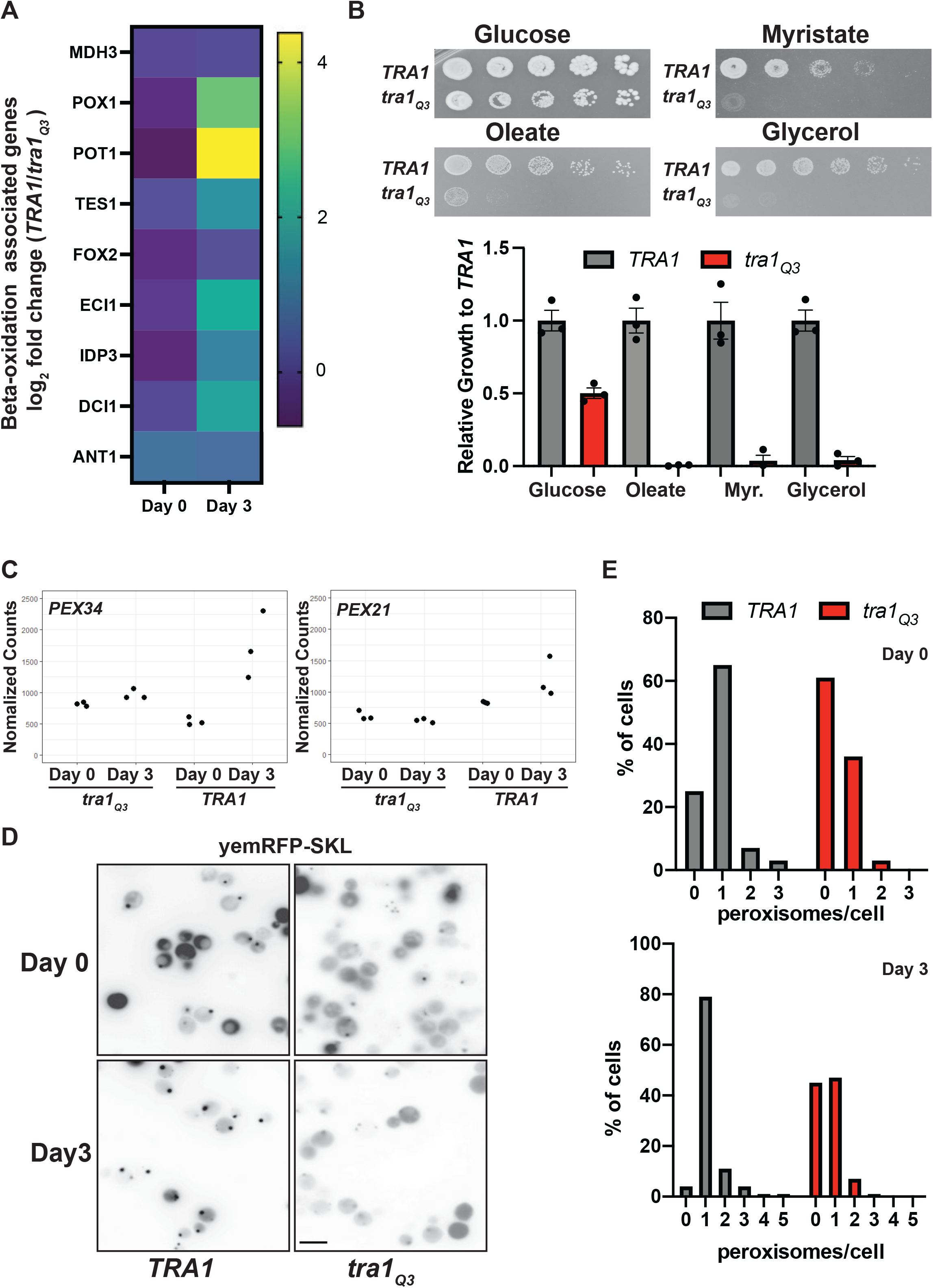
*tra1*_*Q3*_ results in defective peroxisomes. **(A)** log_2_ fold change (*TRA1/tra1*_*Q3*_) for beta-oxidation genes at day 0 and day 3. **(B)** *tra1*_*Q3*_ cells show decreased growth in the presence of oleic acid. *TRA1* and *tra1*_*Q3*_ cells were spotted on agar plates containing either glucose (YPD), oleate, myristate or glycerol as the carbon source. Growth of *tra1*_*Q3*_ cells relative to *TRA1* cells was quantified and is shown in bar graph. **(C)** Normalized RNA sequencing read counts are shown for *PEX34* and *PEX21* in wild-type *TRA1* and *tra1*_*Q3*_ cells at day 0 and day 3. **(D)** *TRA1* and *tra1*_*Q3*_ cells expressing yemRFP-SKL were imaged using fluorescence microscopy at day 0 and day 3 of the aging process. The number of peroxisomes/cell is shown in bar graphs. Bar: 10µm

## DISCUSSION

### A role for Tra1 in chronological aging

Tra1 has been proposed to act as protein-protein hub where it interacts with transcription activators to recruit SAGA and NuA4 co-activator complexes to specific promoters (Elías-Villalobos *et al*. 2019b). In both SAGA and NuA4, Tra1 regulates multiple stress responses associated with the aging process such as cell wall integrity, ethanol sensitivity, protein misfolding, lipid synthesis and the response to DNA damage. Thus, Tra1 regulates various aspects of the transcriptional response to the aging process. Interestingly, the Tra1 homologue in *Drosophila*, Nipped-A, is required to maintain proliferative capacity of intestinal stem cells through aging, indicating that Tra1 may play a role in aging across species (Tauc *et al*. 2017).

This study and previous work using the *tra1*_*Q3*_ mutant (Berg *et al*. 2018; Razzaq *et al*. 2021) support an important regulatory role for the Tra1 PI3K domain despite the absence of key kinase motifs (McMahon *et al*. 1998; Helmlinger *et al*. 2011). Our previous work showed that Tra1_Q3_ has decreased association with SAGA and NuA components (Berg *et al*. 2018). This included Spt20, which is the preferential Tra1 interactor in both fission and budding yeast (Liu *et al*. 2019; Elías-Villalobos *et al*. 2019a; Wang *et al*. 2020). Moreover, Tra1 regulates the incorporation of DUB module components into SAGA (Elías-Villalobos *et al*. 2019a). Interestingly, we found decreased expression of DUB components in aged *tra1*_*Q3*_ cells. Whether this is a response to misassembly of the complex remains to be determined. Similarly, (Leo *et al*. 2018) demonstrated that SAGA, through Gcn5, is essential to properly maintain Ubp8 levels under respiratory conditions. Aging also exacerbated the increase in *TRA1* expression that we previously observed in *tra1*_*Q3*_ cells (Berg *et al*. 2018). The nature of this feedback mechanism remains unclear. Upregulation of *TRA1* is not observed in response to deleting other components of SAGA and NuA4 (Berg *et al*. 2018). *TRA1* expression is also upregulated upon protein misfolding stress (Jiang *et al*. 2019). Therefore, it is reasonable to postulate that increased proteotoxic stress associated with the aging *tra1*_*Q3*_ allele plays a role in this phenotype.

### HAT complexes and lifespan regulation

While we show here that compromising Tra1 function shortens chronological lifespan, the role of the different components of the SAGA and NuA4 complexes in aging is complex. Deleting *GCN5* and *SPT20* shortens chronological lifespan in winemaking yeast (Orozco *et al*. 2012; Picazo *et al*. 2015). Conversely, deleting the SAGA component *SGF11* extends chronological lifespan (Garay *et al*. 2014) indicating that members of HAT complexes can differentially impact the aging process. The SAGA DUB module also regulates replicative aging via its interaction with Sir2 (McCormick *et al*. 2014; Mason *et al*. 2017). Deletion of *GCN5* and its pharmacological inhibition extends replicative lifespan (Huang *et al*. 2020). Therefore, different components of SAGA can differentially affect aging. Arp4p (actin-related protein 4), an essential component of the NuA4 complex, is necessary for chronological aging through its interaction with Hho1 (Miloshev *et al*. 2019). Thus, different components of SAGA and NuA4 can differentially affect aging.

Further studies will be required to determine the role of Tra1 and the impact of the *tra1*_*Q3*_ mutation on the global acetylation of both histone and non-histone substrates. Interestingly, *tra1*_*Q3*_ prevents upregulation of *ACS1* during aging (**Table S2**). *ACS1* ecodes a nucleocytoplasmic acetyl-CoA synthetase whose expression increases during chronological aging (Lesur and Campbell 2004; Wierman *et al*. 2017), presumably to maintain the pool of acetyl-CoA required for histone acetylation (Takahashi *et al*. 2006). *ACS1* null cells consequently display reduced chronological lifespan (Marek and Korona 2013). This is also in agreement with previous observations that cells deleted for NuA4 and SAGA components (*eaf7Δ* and *gcn5Δ*) display decreased levels of acetyl-CoA (Rollins *et al*. 2017).

### Tra1, mitochondria, peroxisomes and the aging process

Here, we showed that several genes important for **β**-oxidation are decreased in *tra1*_*Q3*_ cells after chronological aging. During chronological aging, yeast cells utilize internal fat stores (Goldberg *et al*. 2009). Ultimately, free fatty acids are the substrate for peroxisomal **β**-oxidation. This process allows cells to produce acetyl-CoA that is used to generate the ATP in the mitochondria required for survival at stationary phase. Thus, cells that lack mitochondrial respiration (i.e. incapable of growing on glycerol) cannot grow on oleate (Lockshon *et al*. 2007). In our case, the absence of growth of the *tra1*_*Q3*_ strain on medium containing oleate as the carbon source (**Figure 7b**) could be linked to deficient mitochondrial respiration. In *S. cerevisiae*, Gcn5 and Ubp8 are also required for respiration (Canzonetta *et al*. 2016; Leo *et al*. 2018), a process essential for chronological aging (Pan *et al*. 2011; Ocampo *et al*. 2012).

We previously found that a *TRA1* mutant displays negative genetic interactions with genes associated with mitochondrial function (Hoke *et al*. 2008). This could reflect altered regulation of the retrograde response associated with aging (Kim *et al*. 2004; Jazwinski 2005; Friis *et al*. 2014; Pogoda *et al*. 2021). The mitochondrial retrograde pathway signals to the nucleus via the Rtg proteins (Rtg1, 2 and 3) to upregulate genes associated with mitochondrial stress (Liao and Butow 1993; Ždralević *et al*. 2015). The canonical retrograde target is *CIT2*, which encodes the peroxisomal isoform of citrate synthase. The retrograde pathway regulates expression of several other peroxisomal proteins (Chelstowska and Butow 1995). Retrograde signaling also upregulates other genes involved in the TCA cycle, such as mitochondrial citrate synthase (*CIT1*), aconitase (*ACO1*), and NAD+-dependent isocitrate dehydrogenase (*IDH1*/*2*). Interestingly, we found that these targets of the retrograde pathway are downregulated in aged *tra1*_*Q3*_ (**Table S2**). While there is debate concerning the incorporation of Rtg2 in the SAGA-like (SILK) complex (Pray-Grant *et al*. 2002; Adamus *et al*. 2021), our data suggest a role for Tra1 in the regulation of retrograde signaling that could contribute to the changes in peroxisomal gene expression observed in the aged *tra1*_*Q3*_ cells.

Cells carrying deletions in genes encoding other SAGA components display reduced levels of peroxisomal genes (Ratnakumar and Young 2010) suggesting a further link between HAT complexes and peroxisomal biogenesis/functions. Interestingly, cells incompetent for **β**-oxidation have a less severe aging phenotype than cells devoid of peroxisomes, indicating that other peroxisomal functions are important in regulating chronological lifespan (Lefevre *et al*. 2013). Peroxisome proliferation is also associated with the early stage of replicative aging (Deb *et al*. 2022). Free oxidative radicals have long been suggested to regulate the aging process (Harman 1972). Peroxisomes contain catalase that metabolizes hydrogen peroxide and maintains the cellular redox balance (Lismont *et al*. 2015). Efficient import of catalase into the peroxisomes improves longevity in human cells (Koepke *et al*. 2007). Inhibiting human peroxisomal catalase triggers increased mitochondrial reactive oxygen species (Walton and Pizzitelli 2012). In contrast, deletion and pharmacological inactivation of either form of the yeast catalase (*CTT1*, cytosolic; *CTA*1, peroxisomal), is associated with extended chronological lifespan (Mesquita *et al*. 2010). It was proposed that lack of catalase in young cells triggers a sublethal level of oxidative stress that allows hormetic adaptation to oxidative stress and, consequently, lifespan extension (Mesquita *et al*. 2010). However, overexpressing catalase also extends the chronological lifespan of cells lacking superoxide dismutase (*Δsod1*) showing that catalase levels are indeed important to alleviate oxidative damage associated with aging (Rona *et al*. 2015). Therefore, inability of *tra1*_*Q3*_ cells to properly regulate catalase expression likely has an important role in chronological aging. This is supported by previous studies showing that SAGA plays a major role in the transcriptional response to oxidative stress (Huisinga and Pugh 2004; Sansó *et al*. 2011; Kim *et al*. 2019).

In conclusion, Tra1 regulates gene expression associated with several stress pathways such as regulation of cell wall integrity, mitochondria respiration and peroxisomal function that are crucial for chronological aging.

## ACKNOWLEDGEMENTS

We thank Sonja Di Gregorio for technical assistance with the experiments and Martin Duennwald, Paul Walton and Michael Downey for useful comments along the realization of this project. This study was supported by a Canadian Institutes for Health Research (CIHR) Project Grant (PJT 168882), a National Sciences and Engineering Research Council (NSERC) Discovery Grant (RGPIN-2015-06400) and a John R Evans Leader Fund Grant (65183) to PL. KAB is supported by a Dean’s Research Scholarship from the Schulich Faculty of Medicine & Dentistry at The University of Western Ontario. CJB is supported by an NSERC Discovery Grant (RGPIN-2015-04394). MDB was supported by an NSERC Alexander Graham Bell Canada Graduate Scholarship.

## Figure Legends

**Figure S1:**
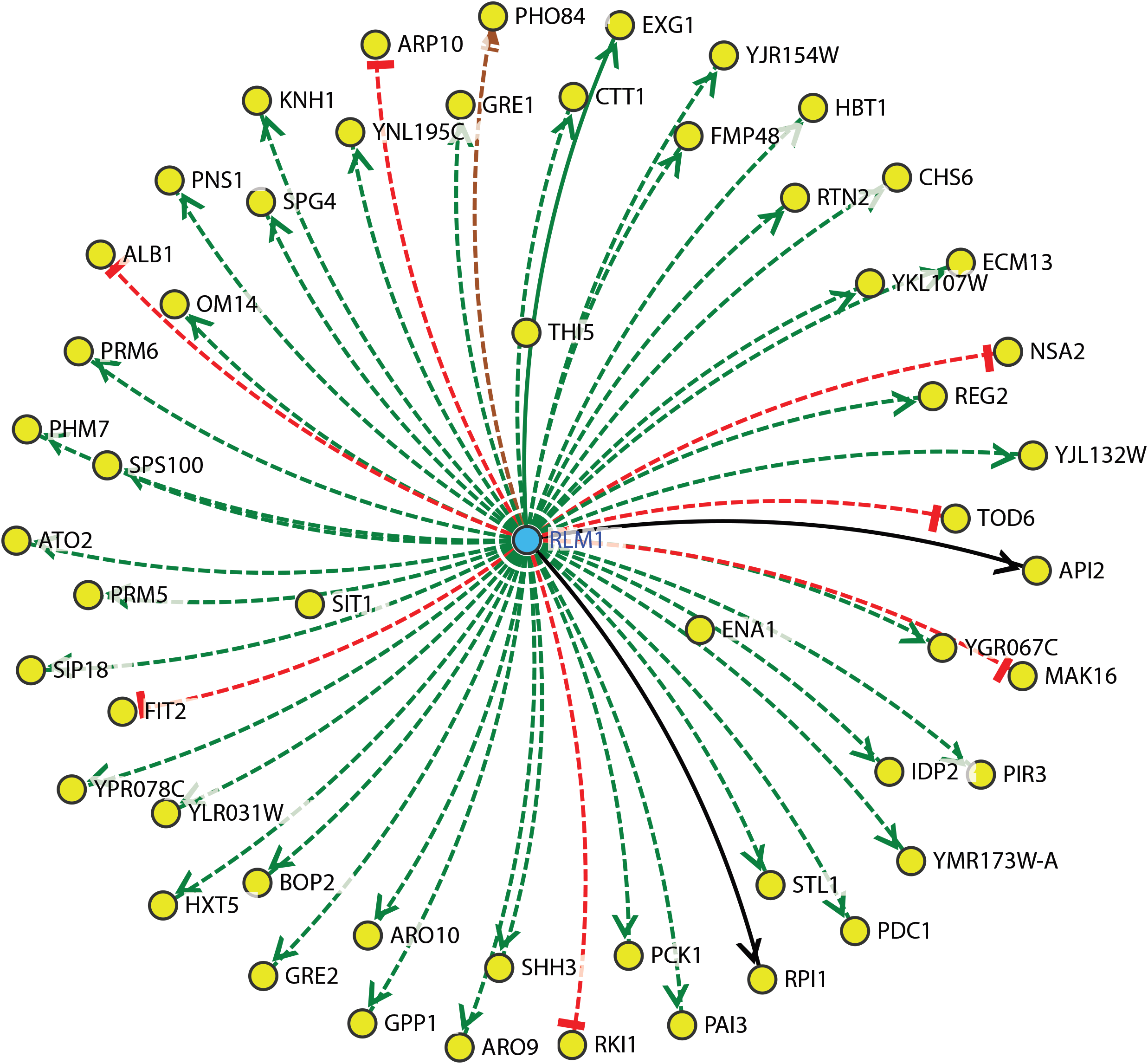
Rlm1 associations with differentially downregulated genes in the aged *tra1*_*Q3*_ strain. The experimental evidence underlying each regulatory association (solid lines for DNA-binding evidence; dashed lines for expression evidence), as well as the sign of the interaction—positive (green), negative (red), positive and negative (brown), or undefined (black) are shown.

**Table S1.**
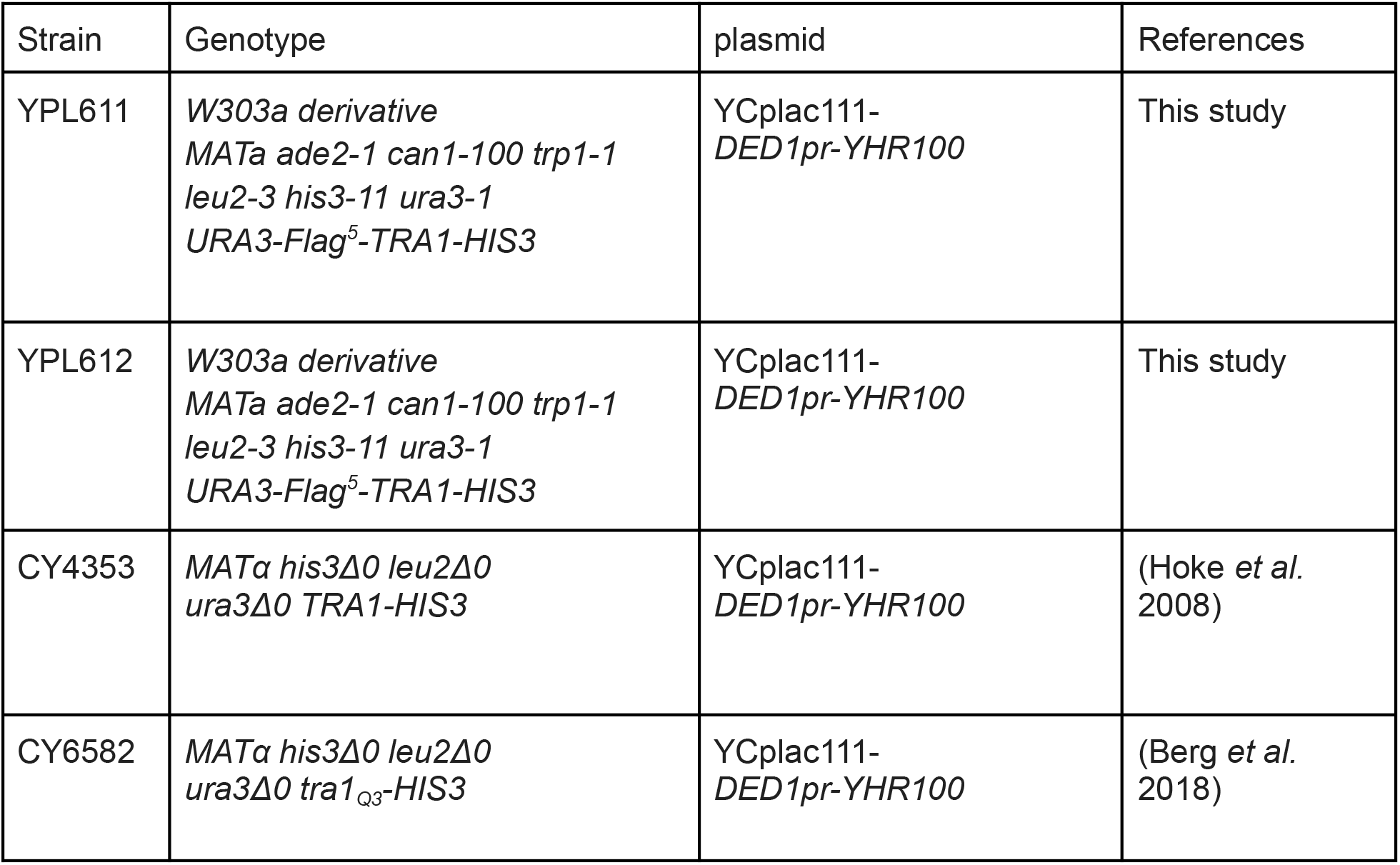

## REFERENCES

Adamus K., C. Reboul, J. Voss, C. Huang, R. B. Schittenhelm, et al., 2021 SAGA and SAGA-like SLIK transcriptional coactivators are structurally and biochemically equivalent. J. Biol. Chem. 296: 100671.

Agarwal S., S. Sharma, V. Agrawal, and N. Roy, 2005 Caloric restriction augments ROS defense in S. cerevisiae, by a Sir2p independent mechanism. Free Radic. Res. 39: 55–62.

Allard S., R. T. Utley, J. Savard, A. Clarke, P. Grant, et al., 1999 NuA4, an essential transcription adaptor/histone H4 acetyltransferase complex containing Esa1p and the ATM-related cofactor Tra1p. EMBO J. 18: 5108–5119.

Arlia-Ciommo A., A. Leonov, A. Piano, V. Svistkova, and V. I. Titorenko, 2014 Cell-autonomous mechanisms of chronological aging in the yeast. Microb. Cell Fact. 1: 163–178.

Atanassov B. S., Y. A. Evrard, A. S. Multani, Z. Zhang, L. Tora, et al., 2009 Gcn5 and SAGA regulate shelterin protein turnover and telomere maintenance. Mol. Cell 35: 352–364.

Beach A., V. R. Richard, A. Leonov, M. T. Burstein, S. D. Bourque, et al., 2013 Mitochondrial membrane lipidome defines yeast longevity. Aging 5: 551–574.

Beach A., A. Leonov, A. Arlia-Ciommo, V. Svistkova, V. Lutchman, et al., 2015 Mechanisms by which different functional states of mitochondria define yeast longevity. Int. J. Mol. Sci. 16: 5528–5554.

Berg M. D., J. Genereaux, J. Karagiannis, and C. J. Brandl, 2018 The Pseudokinase Domain of Tra1 Is Required for Nuclear Localization and Incorporation into the SAGA and NuA4 Complexes. G3 8: 1943–1957.

Bhaumik S. R., T. Raha, D. P. Aiello, and M. R. Green, 2004 In vivo target of a transcriptional activator revealed by fluorescence resonance energy transfer. Genes Dev. 18: 333–343.

Bird A. W., D. Y. Yu, M. G. Pray-Grant, Q. Qiu, K. E. Harmon, et al., 2002 Acetylation of histone H4 by Esa1 is required for DNA double-strand break repair. Nature 419: 411–415.

Bolger A. M., M. Lohse, and B. Usadel, 2014 Trimmomatic: a flexible trimmer for Illumina sequence data. Bioinformatics 30: 2114–2120.

Booth L. N., and A. Brunet, 2016 The Aging Epigenome. Mol. Cell 62: 728–744.

Bosotti R., A. Isacchi, and E. L. Sonnhammer, 2000 FAT: a novel domain in PIK-related kinases. Trends Biochem. Sci. 25: 225–227.

Brown C. E., L. Howe, K. Sousa, S. C. Alley, M. J. Carrozza, et al., 2001 Recruitment of HAT complexes by direct activator interactions with the ATM-related Tra1 subunit. Science 292: 2333–2337.

Burhans W. C., and M. Weinberger, 2009 Acetic acid effects on aging in budding yeast: are they relevant to aging in higher eukaryotes? Cell Cycle 8: 2300–2302.

Burtner C. R., C. J. Murakami, B. K. Kennedy, and M. Kaeberlein, 2009 A molecular mechanism of chronological aging in yeast. Cell Cycle 8: 1256–1270.

Campos S. E., J. A. Avelar-Rivas, E. Garay, A. Juárez-Reyes, and A. DeLuna, 2018 Genomewide mechanisms of chronological longevity by dietary restriction in budding yeast. Aging Cell 17: e12749.

Canzonetta C., M. Leo, S. R. Guarino, A. Montanari, S. Francisci, et al., 2016 SAGA complex and Gcn5 are necessary for respiration in budding yeast. Biochim. Biophys. Acta 1863: 3160–3168.

Chadwick S. R., A. D. Pananos, S. E. Di Gregorio, A. E. Park, P. Etedali-Zadeh, et al., 2016 A toolbox for rapid quantitative assessment of chronological lifespan and survival in Saccharomyces cerevisiae. Traffic 17: 689–703.

Chadwick S. R., and P. Lajoie, 2019 Endoplasmic reticulum stress coping mechanisms and lifespan regulation in health and diseases. Front Cell Dev Biol 7: 84.

Chadwick S. R., E. N. Fazio, P. Etedali-Zadeh, J. Genereaux, M. L. Duennwald, et al., 2020 A functional unfolded protein response is required for chronological aging in Saccharomyces cerevisiae. Curr. Genet. 66: 263–277.

Chaves S. R., A. Rego, V. M. Martins, C. Santos-Pereira, M. J. Sousa, et al., 2021 Regulation of cell death induced by acetic acid in yeasts. Front Cell Dev Biol 9: 642375.

Chelstowska A., and R. A. Butow, 1995 RTG genes in yeast that function in communication between mitochondria and the nucleus are also required for expression of genes encoding peroxisomal proteins. J. Biol. Chem. 270: 18141–18146.

Cheng X., O. Jobin-Robitaille, P. Billon, R. Buisson, H. Niu, et al., 2018 Phospho-dependent recruitment of the yeast NuA4 acetyltransferase complex by MRX at DNA breaks regulates RPA dynamics during resection. Proceedings of the National Academy of Sciences 115: 10028–10033.

Cheng X., V. Côté, and J. Côté, 2021 NuA4 and SAGA acetyltransferase complexes cooperate for repair of DNA breaks by homologous recombination. PLoS Genet. 17: e1009459.

Cheung A. C. M., and L. M. Díaz-Santín, 2019 Share and share alike: the role of Tra1 from the SAGA and NuA4 coactivator complexes. Transcription 10: 37–43.

Choi K.-M., Y.-Y. Kwon, and C.-K. Lee, 2013 Characterization of global gene expression during assurance of lifespan extension by caloric restriction in budding yeast. Exp. Gerontol. 48: 1455–1468.

Clarke A. S., J. E. Lowell, S. J. Jacobson, and L. Pillus, 1999 Esa1p is an essential histone acetyltransferase required for cell cycle progression. Mol. Cell. Biol. 19: 2515–2526.

Deb R., S. Ghose, and S. Nagotu, 2022 Increased peroxisome proliferation is associated with early yeast replicative ageing. Curr. Genet. 68: 207–225.

De Luca V., M. Leo, E. Cretella, A. Montanari, M. Saliola, et al., 2022 Role of yUbp8 in mitochondria and hypoxia entangles the finding of human ortholog usp22 in the glioblastoma pseudo-palisade microlayer. Cells 11. https://doi.org/10.3390/cells11101682

Deprez M.-A., E. Eskes, T. Wilms, P. Ludovico, and J. Winderickx, 2018 pH homeostasis links the nutrient sensing PKA/TORC1/Sch9 ménage-à-trois to stress tolerance and longevity. Microb. Cell Fact. 5: 119–136.

Díaz-Santín L. M., N. Lukoyanova, E. Aciyan, and A. C. M. Cheung, 2017 Cryo-EM structure of the SAGA and NuA4 coactivator subunit Tra1 at 3.7 angstrom resolution. https://doi.org/10.7554/eLife.28384

Dobin A., C. A. Davis, F. Schlesinger, J. Drenkow, C. Zaleski, et al., 2013 STAR: ultrafast universal RNA-seq aligner. Bioinformatics 29: 15–21.

Downey M., 2021 Non-histone protein acetylation by the evolutionarily conserved GCN5 and PCAF acetyltransferases. Biochim. Biophys. Acta Gene Regul. Mech. 1864: 194608.

Downs J. A., S. Allard, O. Jobin-Robitaille, A. Javaheri, A. Auger, et al., 2004 Binding of chromatin-modifying activities to phosphorylated histone H2A at DNA damage sites. Mol. Cell 16: 979–990.

Eberharter A., and P. B. Becker, 2002 Histone acetylation: a switch between repressive and permissive chromatin. Second in review series on chromatin dynamics. EMBO Rep. 3. https://doi.org/10.1093/embo-reports/kvf053

Edgar R., M. Domrachev, and A. E. Lash, 2002 Gene Expression Omnibus: NCBI gene expression and hybridization array data repository. Nucleic Acids Res. 30: 207–210.

Elías-Villalobos A., D. Toullec, C. Faux, M. Séveno, and D. Helmlinger, 2019a Chaperone-mediated ordered assembly of the SAGA and NuA4 transcription co-activator complexes in yeast. Nat. Commun. 10: 5237.

Elías-Villalobos A., P. Fort, and D. Helmlinger, 2019b New insights into the evolutionary conservation of the sole PIKK pseudokinase Tra1/TRRAP. Biochem. Soc. Trans. 47: 1597–1608.

Fabrizio P., F. Pozza, S. D. Pletcher, C. M. Gendron, and V. D. Longo, 2001 Regulation of longevity and stress resistance by Sch9 in yeast. Science 292: 288–290.

Fontana L., L. Partridge, and V. D. Longo, 2010 Extending healthy life span—from yeast to humans. Science 328: 321–326.

Friis R. M. N., J. P. Glaves, T. Huan, L. Li, B. D. Sykes, et al., 2014 Rewiring AMPK and mitochondrial retrograde signaling for metabolic control of aging and histone acetylation in respiratory-defective cells. Cell Rep. 7: 565–574.

Garay E., S. E. Campos, J. González de la Cruz, A. P. Gaspar, A. Jinich, et al., 2014 High-resolution profiling of stationary-phase survival reveals yeast longevity factors and their genetic interactions. PLoS Genet. 10: e1004168.

Ghavidel A., K. Baxi, M. Prusinkiewicz, C. Swan, Z. R. Belak, et al., 2018 Rapid nuclear exclusion of Hcm1 in aging leads to vacuolar alkalization and replicative senescence. G3 8: 1579–1592.

Goldberg A. A., S. D. Bourque, P. Kyryakov, C. Gregg, T. Boukh-Viner, et al., 2009 Effect of calorie restriction on the metabolic history of chronologically aging yeast. Exp. Gerontol. 44: 555–571.

Grant P. A., L. Duggan, J. Côté, S. M. Roberts, J. E. Brownell, et al., 1997 Yeast Gcn5 functions in two multisubunit complexes to acetylate nucleosomal histones: characterization of an Ada complex and the SAGA (Spt/Ada) complex. Genes Dev. 11: 1640–1650.

Grant P. A., D. Schieltz, M. G. Pray-Grant, J. R. Yates 3rd, and J. L. Workman, 1998 The ATM-related cofactor Tra1 is a component of the purified SAGA complex. Mol. Cell 2: 863–867.

Guaragnella N., M. Stirpe, D. Marzulli, C. Mazzoni, and S. Giannattasio, 2019 Acid stress triggers resistance to acetic acid-induced regulated cell death through activation which requires RTG2 in Yeast. Oxid. Med. Cell. Longev. 2019: 4651062.

Harman D., 1972 The biologic clock: the mitochondria? J. Am. Geriatr. Soc. 20: 145–147.

Helmlinger D., S. Marguerat, J. Villén, D. L. Swaney, S. P. Gygi, et al., 2011 Tra1 has specific regulatory roles, rather than global functions, within the SAGA co-activator complex. EMBO J. 30: 2843–2852.

Henry K. W., A. Wyce, W.-S. Lo, L. J. Duggan, N. C. Tolga Emre, et al., 2003 Transcriptional activation via sequential histone H2B ubiquitylation and deubiquitylation, mediated by SAGA-associated Ubp8. Genes Dev. 17: 2648–2663.

Hill R., and P. W. K. Lee, 2010 The DNA-dependent protein kinase (DNA-PK): More than just a case of making ends meet? Cell Cycle 9: 3460–3469.

Hiltunen J. K., A. M. Mursula, H. Rottensteiner, R. K. Wierenga, A. J. Kastaniotis, et al., 2003 The biochemistry of peroxisomal beta-oxidation in the yeast Saccharomyces cerevisiae. FEMS Microbiol. Rev. 27: 35–64.

Hoke S. M. T., J. Guzzo, B. Andrews, and C. J. Brandl, 2008 Systematic genetic array analysis links the Saccharomyces cerevisiae SAGA/SLIK and NuA4 component Tra1 to multiple cellular processes. BMC Genet. 9: 46.

Hu J., M. Wei, H. Mirzaei, F. Madia, M. Mirisola, et al., 2014 Tor-Sch9 deficiency activates catabolism of the ketone body-like acetic acid to promote trehalose accumulation and longevity. Aging Cell 13: 457–467.

Huang B., D. Zhong, J. Zhu, Y. An, M. Gao, et al., 2020 Inhibition of histone acetyltransferase GCN5 extends lifespan in both yeast and human cell lines. Aging Cell 19: e13129.

Huisinga K. L., and B. F. Pugh, 2004 A genome-wide housekeeping role for TFIID and a highly regulated stress-related role for SAGA in Saccharomyces cerevisiae. Mol. Cell 13: 573–585.

Huse M., and J. Kuriyan, 2002 The conformational plasticity of protein kinases. Cell 109: 275–282.

Jazwinski S. M., 2005 The retrograde response links metabolism with stress responses, chromatin-dependent gene activation, and genome stability in yeast aging. Gene 354: 22–27.

Jiang J. C., E. Jaruga, M. V. Repnevskaya, and S. M. Jazwinski, 2000 An intervention resembling caloric restriction prolongs life span and retards aging in yeast. FASEB J. 14: 2135–2137.

Jiang Y., M. D. Berg, J. Genereaux, K. Ahmed, M. L. Duennwald, et al., 2019 Sfp1 links TORC1 and cell growth regulation to the yeast SAGA-complex component Tra1 in response to polyQ proteotoxicity. Traffic 20: 267–283.

Kaeberlein M., M. McVey, and L. Guarente, 1999 The SIR2/3/4 complex and SIR2 alone promote longevity in Saccharomyces cerevisiae by two different mechanisms. Genes Dev. 13: 2570–2580.

Kaeberlein M., 2010 Lessons on longevity from budding yeast. Nature 464: 513–519.

Kane A. E., and D. A. Sinclair, 2019 Epigenetic changes during aging and their reprogramming potential. Crit. Rev. Biochem. Mol. Biol. 54: 61–83.

Keith C. T., and S. L. Schreiber, 1995 PIK-related kinases: DNA repair, recombination, and cell cycle checkpoints. Science 270: 50–51.

Kim S., K. Ohkuni, E. Couplan, and S. M. Jazwinski, 2004 The histone acetyltransferase GCN5 modulates the retrograde response and genome stability determining yeast longevity. Biogerontology 5: 305–316.

Kim M., Y. Choi, H. Kim, and D. Lee, 2019 SAGA DUBm-mediated surveillance regulates prompt export of stress-inducible transcripts for proteostasis. Nat. Commun. 10: 2458.

Kingston R. E., C. A. Bunker, and A. N. Imbalzano, 1996 Repression and activation by multiprotein complexes that alter chromatin structure. Genes Dev. 10. https://doi.org/10.1101/gad.10.8.905

Knutson B. A., and S. Hahn, 2011 Domains of Tra1 important for activator recruitment and transcription coactivator functions of SAGA and NuA4 complexes. Mol. Cell. Biol. 31: 818–831.

Koepke J. I., K.-A. Nakrieko, C. S. Wood, K. K. Boucher, L. J. Terlecky, et al., 2007 Restoration of peroxisomal catalase import in a model of human cellular aging. Traffic 8: 1590–1600.

Laun P., M. Rinnerthaler, E. Bogengruber, G. Heeren, and M. Breitenbach, 2006 Yeast as a model for chronological and reproductive aging - a comparison. Exp. Gerontol. 41: 1208–1212.

Lefevre S. D., C. W. van Roermund, R. J. A. Wanders, M. Veenhuis, and I. J. van der Klei, 2013 The significance of peroxisome function in chronological aging of Saccharomyces cerevisiae. Aging Cell 12: 784–793.

Leo M., G. Fanelli, S. Di Vito, B. Traversetti, M. La Greca, et al., 2018 Ubiquitin protease Ubp8 is necessary for S. cerevisiae respiration. Biochim. Biophys. Acta Mol. Cell Res. https://doi.org/10.1016/j.bbamcr.2018.07.025

Leonov A., R. Feldman, A. Piano, A. Arlia-Ciommo, V. Lutchman, et al., 2017 Caloric restriction extends yeast chronological lifespan via a mechanism linking cellular aging to cell cycle regulation, maintenance of a quiescent state, entry into a non-quiescent state and survival in the non-quiescent state. Oncotarget 8: 69328–69350.

Lesur I., and J. L. Campbell, 2004 The transcriptome of prematurely aging yeast cells is similar to that of telomerase-deficient cells. Mol. Biol. Cell 15: 1297–1312.

Liao X., and R. A. Butow, 1993 RTG1 and RTG2: two yeast genes required for a novel path of communication from mitochondria to the nucleus. Cell 72: 61–71.

Liao Y., G. K. Smyth, and W. Shi, 2014 featureCounts: an efficient general purpose program for assigning sequence reads to genomic features. Bioinformatics 30: 923–930.

Lin Y.-Y., Y. Qi, J.-Y. Lu, X. Pan, D. S. Yuan, et al., 2008 A comprehensive synthetic genetic interaction network governing yeast histone acetylation and deacetylation. Genes & Development 22: 2062–2074.

Lin L., L. Chamberlain, L. J. Zhu, and M. R. Green, 2012 Analysis of Gal4-directed transcription activation using Tra1 mutants selectively defective for interaction with Gal4. Proc. Natl. Acad. Sci. U. S. A. 109: 1997–2002.

Lismont C., M. Nordgren, P. P. Van Veldhoven, and M. Fransen, 2015 Redox interplay between mitochondria and peroxisomes. Front Cell Dev Biol 3: 35.

Liu G., X. Zheng, H. Guan, Y. Cao, H. Qu, et al., 2019 Architecture of Saccharomyces cerevisiae SAGA complex. Cell Discovery 5.

Lockshon D., L. E. Surface, E. O. Kerr, M. Kaeberlein, and B. K. Kennedy, 2007 The sensitivity of yeast mutants to oleic acid implicates the peroxisome and other processes in membrane function. Genetics 175: 77–91.

Longo V. D., 1999 Mutations in signal transduction proteins increase stress resistance and longevity in yeast, nematodes, fruit flies, and mammalian neuronal cells. Neurobiol. Aging 20: 479–486.

Longo V. D., and P. Fabrizio, 2012 Chronological aging in Saccharomyces cerevisiae. Subcell. Biochem. 57: 101–121.

Love M. I., W. Huber, and S. Anders, 2014 Moderated estimation of fold change and dispersion for RNA-seq data with DESeq2. Genome Biol. 15: 550.

MacLean M., N. Harris, and P. W. Piper, 2001 Chronological lifespan of stationary phase yeast cells; a model for investigating the factors that might influence the ageing of postmitotic tissues in higher organisms. Yeast 18: 499–509.

Marek A., and R. Korona, 2013 Restricted pleiotropy facilitates mutational erosion of major life-history traits. Evolution 67: 3077–3086.

Mason A. G., R. M. Garza, M. A. McCormick, B. Patel, B. K. Kennedy, et al., 2017 The replicative lifespan-extending deletion of SGF73 results in altered ribosomal gene expression in yeast. Aging Cell 16: 785–796.

Matecic M., D. L. Smith, X. Pan, N. Maqani, S. Bekiranov, et al., 2010 A microarray-based genetic screen for yeast chronological aging factors. PLoS Genet. 6: e1000921.

McCormick M. A., A. G. Mason, S. J. Guyenet, W. Dang, R. M. Garza, et al., 2014 The SAGA histone deubiquitinase module controls yeast replicative lifespan via Sir2 interaction. Cell Rep. 8: 477–486.

McMahon S. B., H. A. Van Buskirk, K. A. Dugan, T. D. Copeland, and M. D. Cole, 1998 The novel ATM-related protein TRRAP is an essential cofactor for the c-Myc and E2F oncoproteins. Cell 94: 363–374.

Medkour Y., and V. I. Titorenko, 2016 Mitochondria operate as signaling platforms in yeast aging. Aging 8: 212–213.

Mesquita A., M. Weinberger, A. Silva, B. Sampaio-Marques, B. Almeida, et al., 2010 Caloric restriction or catalase inactivation extends yeast chronological lifespan by inducing H2O2 and superoxide dismutase activity. Proc. Natl. Acad. Sci. U. S. A. 107: 15123–15128.

Miloshev G., D. Staneva, K. Uzunova, B. Vasileva, M. Draganova-Filipova, et al., 2019 Linker histones and chromatin remodelling complexes maintain genome stability and control cellular ageing. Mechanisms of Ageing and Development 177: 55–65.

Mirisola M. G., and V. D. Longo, 2022 Yeast chronological lifespan: longevity regulatory genes and mechanisms. Cells 11. https://doi.org/10.3390/cells11101714

Mohammad K., J. A. Baratang Junio, T. Tafakori, E. Orfanos, and V. I. Titorenko, 2020 Mechanisms that link chronological aging to cellular quiescence in budding yeast. Int. J. Mol. Sci. 21. https://doi.org/10.3390/ijms21134717

Monteiro P. T., J. Oliveira, P. Pais, M. Antunes, M. Palma, et al., 2020 YEASTRACT : a portal for cross-species comparative genomics of transcription regulation in yeasts. Nucleic Acids Research 48: D642–D649.

Mordes D. A., G. G. Glick, R. Zhao, and D. Cortez, 2008 TopBP1 activates ATR through ATRIP and a PIKK regulatory domain. Genes & Development 22: 1478–1489.

Morgan M. T., M. Haj-Yahya, A. E. Ringel, P. Bandi, A. Brik, et al., 2016 Structural basis for histone H2B deubiquitination by the SAGA DUB module. Science 351: 725–728.

Mortimer R. K., and J. R. Johnston, 1959 Life span of individual yeast cells. Nature 183: 1751–1752.

Mumberg D., R. Müller, and M. Funk, 1995 Yeast vectors for the controlled expression of heterologous proteins in different genetic backgrounds. Gene 156: 119–122.

Murakami C. J., C. R. Burtner, B. K. Kennedy, and M. Kaeberlein, 2008 A method for high-throughput quantitative analysis of yeast chronological life span. J. Gerontol. A Biol. Sci. Med. Sci. 63: 113–121.

Mutiu A. I., S. M. T. Hoke, J. Genereaux, C. Hannam, K. MacKenzie, et al., 2007 Structure/function analysis of the phosphatidylinositol-3-kinase domain of yeast tra1. Genetics 177: 151–166.

Nelson J. D., O. Denisenko, and K. Bomsztyk, 2006 Protocol for the fast chromatin immunoprecipitation (ChIP) method. Nat. Protoc. 1: 179–185.

Ning Z., X. Guo, X. Liu, C. Lu, A. Wang, et al., 2022 USP22 regulates lipidome accumulation by stabilizing PPARγ in hepatocellular carcinoma. Nat. Commun. 13: 2187.

Ocampo A., J. Liu, E. A. Schroeder, G. S. Shadel, and A. Barrientos, 2012 Mitochondrial respiratory thresholds regulate yeast chronological life span and its extension by caloric restriction. Cell Metab. 16: 55–67.

Orozco H., E. Matallana, and A. Aranda, 2012 Wine yeast sirtuins and Gcn5p control aging and metabolism in a natural growth medium. Mech. Ageing Dev. 133: 348–358.

Pan Y., E. A. Schroeder, A. Ocampo, A. Barrientos, and G. S. Shadel, 2011 Regulation of yeast chronological life span by TORC1 via adaptive mitochondrial ROS signaling. Cell Metab. 13: 668–678.

Pan Y., 2011 Mitochondria, reactive oxygen species, and chronological aging: a message from yeast. Exp. Gerontol. 46: 847–852.

Pavletich N. P., and H. Yang, 2013 mTOR kinase structure, mechanism and regulation

Petriv O. I., and R. A. Rachubinski, 2004 Lack of peroxisomal catalase causes a progeric phenotype in Caenorhabditis elegans. J. Biol. Chem. 279: 19996–20001.

Petropavlovskiy A. A., M. G. Tauro, P. Lajoie, and M. L. Duennwald, 2020 A quantitative imaging-based protocol for yeast growth and survival on agar plates. STAR Protoc 1: 100182.

Petrova V. Y., D. Drescher, A. V. Kujumdzieva, and M. J. Schmitt, 2004 Dual targeting of yeast catalase A to peroxisomes and mitochondria. Biochem. J 380: 393–400.

Picazo C., H. Orozco, E. Matallana, and A. Aranda, 2015 Interplay among Gcn5, Sch9 and mitochondria during chronological aging of wine yeast is dependent on growth conditions. PLoS One 10: e0117267.

Pogoda E., H. Tutaj, A. Pirog, K. Tomala, and R. Korona, 2021 Overexpression of a single ORF can extend chronological lifespan in yeast if retrograde signaling and stress response are stimulated. Biogerontology 22: 415–427.

Pray-Grant M. G., D. Schieltz, S. J. McMahon, J. M. Wood, E. L. Kennedy, et al., 2002 The novel SLIK histone acetyltransferase complex functions in the yeast retrograde response pathway. Mol. Cell. Biol. 22: 8774–8786.

Prokakis E., A. Dyas, R. Grün, S. Fritzsche, U. Bedi, et al., 2021 USP22 promotes HER2-driven mammary carcinoma aggressiveness by suppressing the unfolded protein response. Oncogene 40: 4004–4018.

Purdue P. E., X. Yang, and P. B. Lazarow, 1998 Pex18p and Pex21p, a novel pair of related peroxins essential for peroxisomal targeting by the PTS2 pathway. J. Cell Biol. 143: 1859–1869.

Rando O. J., and F. Winston, 2012 Chromatin and transcription in yeast. Genetics 190: 351–387.

Ratnakumar S., and E. T. Young, 2010 Snf1 dependence of peroxisomal gene expression is mediated by Adr1. J. Biol. Chem. 285: 10703–10714.

Razzaq I., M. D. Berg, Y. Jiang, J. Genereaux, D. Uthayakumar, et al., 2021 The SAGA and NuA4 component Tra1 regulates Candida albicans drug resistance and pathogenesis. Genetics 219. https://doi.org/10.1093/genetics/iyab131

Roberts S. M., and F. Winston, 1997 Essential functional interactions of SAGA, a Saccharomyces cerevisiae complex of Spt, Ada, and Gcn5 proteins, with the Snf/Swi and Srb/Mediator complexes. Genetics 147: 451–465.

Rollins M., S. Huard, A. Morettin, J. Takuski, T. T. Pham, et al., 2017 Lysine acetyltransferase NuA4 and acetyl-CoA regulate glucose-deprived stress granule formation in Saccharomyces cerevisiae. PLoS Genet. 13: e1006626.

Rona G., R. Herdeiro, C. J. Mathias, F. A. Torres, M. D. Pereira, et al., 2015 CTT1 overexpression increases life span of calorie-restricted Saccharomyces cerevisiae deficient in Sod1. Biogerontology 16: 343–351.

Saleh A., D. Schieltz, N. Ting, S. B. McMahon, D. W. Litchfield, et al., 1998 Tra1p is a component of the yeast Ada.Spt transcriptional regulatory complexes. J. Biol. Chem. 273: 26559–26565.

Sansó M., I. Vargas-Pérez, L. Quintales, F. Antequera, J. Ayté, et al., 2011 Gcn5 facilitates Pol II progression, rather than recruitment to nucleosome-depleted stress promoters, in Schizosaccharomyces pombe. Nucleic Acids Res. 39: 6369–6379.

Sardi M., and A. P. Gasch, 2018 Genetic background effects in quantitative genetics: gene-by-system interactions. Curr. Genet. 64: 1173–1176.

Schneider C. A., W. S. Rasband, and K. W. Eliceiri, 2012 NIH Image to ImageJ: 25 years of image analysis. Nat. Methods 9: 671–675.

Seah T. C., and J. G. Kaplan, 1973 Purification and properties of the catalase of bakers’ yeast. J. Biol. Chem. 248: 2889–2893.

Sharov G., K. Voltz, A. Durand, O. Kolesnikova, G. Papai, et al., 2017 Structure of the transcription activator target Tra1 within the chromatin modifying complex SAGA. Nat. Commun. 8: 1556.

Shiloh Y., and Y. Ziv, 2013 The ATM protein kinase: regulating the cellular response to genotoxic stress, and more. Nat. Rev. Mol. Cell Biol. 14: 197–210.

Smith G. C. M., and S. P. Jackson, 2003 The PIKK family of protein kinases. Handbook of Cell Signaling 557–561.

Smith E. N., and L. Kruglyak, 2008 Gene-environment interaction in yeast gene expression. PLoS Biol. 6: e83.

Steinkraus K. A., M. Kaeberlein, and B. K. Kennedy, 2008 Replicative aging in yeast: the means to the end. Annu. Rev. Cell Dev. Biol. 24: 29–54.

Steunou A.-L., D. Rossetto, and J. Côté, 2014 Regulating chromatin by histone acetylation. Fundamentals of Chromatin 147–212.

Takahashi H., J. M. McCaffery, R. A. Irizarry, and J. D. Boeke, 2006 Nucleocytosolic acetyl-coenzyme a synthetase is required for histone acetylation and global transcription. Mol. Cell 23: 207–217.

Tauc H. M., A. Tasdogan, P. Meyer, and P. Pandur, 2017 Nipped-A regulates intestinal stem cell proliferation in. Development 144: 612–623.

Taylor S. S., and A. P. Kornev, 2011 Protein kinases: evolution of dynamic regulatory proteins. Trends in Biochemical Sciences 36: 65–77.

Tissenbaum H. A., and L. Guarente, 2002 Model organisms as a guide to mammalian aging. Dev. Cell 2: 9–19.

Tower R. J., A. Fagarasanu, J. D. Aitchison, and R. A. Rachubinski, 2011 The peroxin Pex34p functions with the Pex11 family of peroxisomal divisional proteins to regulate the peroxisome population in yeast. Mol. Biol. Cell 22: 1727–1738.

Vasileva B., D. Staneva, N. Krasteva, G. Miloshev, and M. Georgieva, 2021 Changes in chromatin organization eradicate cellular stress resilience to UVA/B light and induce premature aging. Cells 10. https://doi.org/10.3390/cells10071755

Walton P. A., and M. Pizzitelli, 2012 Effects of peroxisomal catalase inhibition on mitochondrial function. Front. Physiol. 3: 108.

Wang H., C. Dienemann, A. Stützer, H. Urlaub, A. C. M. Cheung, et al., 2020 Structure of the transcription coactivator SAGA. Nature 577: 717–720.

Wierman M. B., N. Maqani, E. Strickler, M. Li, and J. S. Smith, 2017 Caloric restriction extends yeast chronological life span by optimizing the Snf1 (AMPK) signaling pathway. Mol. Cell. Biol. 37. https://doi.org/10.1128/MCB.00562-16

Yamauchi Y., and S. Izawa, 2016 Prioritized Expression of BTN2 of Saccharomyces cerevisiae under Pronounced Translation Repression Induced by Severe Ethanol Stress. Front. Microbiol. 7: 1319.

Ždralević M., N. Guaragnella, and S. Giannattasio, 2015 Yeast as a tool to study mitochondrial retrograde pathway en route to cell stress response. Methods Mol. Biol. 1265: 321–331.

